# AxoDen: An algorithm for the automated quantification of axonal density in defined brain regions

**DOI:** 10.1101/2024.05.30.596687

**Authors:** Raquel Adaia Sandoval Ortega, Emmy Li, Oliver Joseph, Pascal A. Dufour, Gregory Corder

**Affiliations:** Department of Psychiatry, Perelman School of Medicine, University of Pennsylvania, Philadelphia, PA, USA; department of Neuroscience, Mahoney Institute for Neurosciences, Perelman School of Medicine, University of Pennsylvania, Philadelphia, PA, USA; Department of Anesthesiology and Critical Care, Perelman School of Medicine, University of Pennsylvania, Philadelphia,PA, USA

## Abstract

The rodent brain contains 70,000,000+ neurons interconnected via complex axonal circuits with varying architectures. Neural pathologies are often associated with anatomical changes in these axonal projections and synaptic connections. Notably, axonal density variations of local and long-range projections increase or decrease as a function of the strengthening or weakening, respectively, of the information flow between brain regions. Traditionally, histological quantification of axonal inputs relied on assessing the mean fluorescence intensity within a rectangle placed in the brain region-of-interest. Despite yielding valuable insights, this conventional method is notably susceptible to background fluorescence, post-acquisition adjustments, and inter-researcher variability. Additionally, it fails to account for the non-uniform innervation across brain regions, thus overlooking critical data such as innervation percentages and axonal distribution patterns. In response to these challenges, we introduce AxoDen, an open-source semi-automated platform designed to increase the speed and rigor of axon quantifications for basic neuroscience discovery. AxoDen processes user-defined brain regions-of-interests incorporating dynamic thresholding of grayscales-transformed images to facilitate binarized pixel measurements. Thereby AxoDen segregates the image content into signal and non-signal categories, effectively eliminating background interference and enabling the exclusive measurement of fluorescence from axonal projections. AxoDen provides detailed and accurate representations of axonal density and spatial distribution. AxoDen’s advanced yet user-friendly platform enhances the reliability and efficiency of axonal density analysis and facilitates access to unbiased high-quality data analysis with no technical background or coding experience required. AxoDen is freely available to everyone as a valuable neuroscience tool for dissecting axonal innervation patterns in precisely defined brain regions.

## Introduction

Understanding the brain’s structural integrity and connectivity is fundamental in neuroscience (Luo, 2021). Studies ranging from basic anatomy to complex neurological disorders rely on quantification of axonal projections across brain regions to assess changes in information-processing pathways. Accurate measures of axonal density allow to infer potential mechanisms underlying alterations on cognitive, sensory, and motor functions. Consequently, precise axonal quantification is crucial to reveal the structural connectivity in normal and pathological states.

Traditionally, assessing axonal inputs to specific brain regions-of-interest (ROI) relied on mean fluorescence intensity measurements within a rectangle placed on ROls. This method has shed light on complex connectivity patterns and pathologies effects on neural networks. However, significant limitations including susceptibility to background fluorescence, vulnerability to post-acquisition image adjustments, and the inherent inter-researcher variability, raise concerns regarding reliability and precision. Furthermore, this approach does not accommodate the innervation heterogeneity within ROls, neglecting essential information on spatial distribution, and innervation percentage.

Efforts to automate quantification have led to the development of several tools, each with strengths and limitations. MeDUsA (Nitta et al., 2023), created in Python, uses advanced convolutional neural networks (CNN) to identify an ROI and quantify axon terminals in Drosophila’s visual system. MeDUsA achieves high accuracy for Drosophila, but its high specificity limits its applicability to other species. An algorithm with higher potential for interspecies utilization is AxonTracer (Patel et al., 2018), created as an lmageJ plugin to measure axonal length in rat spinal cord. While this tool yields valuable insights into axonal complexity, a skeletonization process that reduces axons to a uniform width of one pixel prevents quantification of innervation percentages. The lmageJ macro DEFiNE (Powell et al., 2019) advances the methodology by incorporating an automatic pre-processing stage to diminish background fluorescence before semi-automated quantification. DEFiNE evaluates z-stack images eliminating artifacts and defines signal as pixels exceeding 4 standard deviations above the mean intensity of axon-free areas. However, DEFiNE requires dual-channel imaging-one for the target fluorophore and another for background fluorescence-limiting its use in experiments using all channels for multiple information levels. Additionally, DEFiNE’s need for rectangular images precludes comprehensive brain region innervation analysis. Lastly, TrailMap (Friedmann et al., 2019) transcends the limitations of 2D analysis by employing CNNs to map axonal projections within three-dimensional structures. This Python-based algorithm facilitates region-specific quantification of total axonal content in 3D, necessitating intact, cleared mouse brains imaged with light-sheet microscopy. Despite its precision and analysis depth, TrailMap’s reliance on advanced imaging techniques and computational resources places it beyond reach of many research laboratories.

These tools exemplify the evolving landscape of neural imaging and analysis, highlighting the ongoing need for methodologies balancing specificity, versatility, and accessibility to accommodate the diverse requirements of neuroscience research. Simple and reproducible algorithms, such as Axoden, have the benefit of easy interpretability, which is of special importance when used in scientific analysis. This stands in stark contrast to CNNs, which might yield overall more accurate outputs, but can introduce new biases and errors, difficult to understand due to the black-box-behavior of such methods.

## Results

### AxoDen: A New Method for Axonal Density Quantifica-tion

Recognizing the aforementioned challenges, we have built AxoDen **(Fig. lA)** – a versatile histology image analysis platform that refines and simplifies the process of quantifying axonal inputs, is applicable to any animal species and fluorophores, does not require an advanced setup nor GPU, and uses only one channel, or fluorophore. Beyond the traditional reliance on rectangular measurement areas, AxoDen processes images that have been masked and cropped to specifically fit the brain ROls, conforming to the actual contours identified in the brain atlas selected by the user. This approach allows for the isolation and exclusive measurement of fluorescence signals from axonal projections within the defined brain region.

By incorporating dynamic thresholding for the binarization of grayscale images, our protocol distinguishes between signal and non-signal elements within the brain region, effectively minimizing background interference and consequently ensuring data collection focuses solely on axonal information. The quantification of fluorescently labeled axons is performed for the whole ROI and is also projected to both the X and the Y axis for the spatial analysis of projections. The data on the signal percent, ROls area, and threshold used for binarization are saved as data frames that the user can retrieve for later statistical evaluation. Furthermore, AxoDen generates a summary figure containing information on these parameters for the immediate visualization of the results (as mean ± SEM) after running the analysis.

In order to validate AxoDen and compare it to the traditional methods of measuring fluorescence intensity, we used a dataset of 46 images of pain active axons originating from the anterior cingulate cortex (ACC) (James et al., 2024). Two experimenters selected 19 brain ROls from 4 mice and each of them generated a new dataset consisting of 76 items. We first compared the time needed to prepare the images for AxoDen analysis with the time required with the traditional method in which the experimenter collects the mean fluorescence intensity within a rectangular shape placed on the brain ROI. We refer the later as Mean Intensity from now on. Preparing the data for AxoDen was 1.6 times faster than collecting the data for Mean Intensity analysis (AxoDen: 9.68 ± 0.33 hours; Mean Intensity: 15.22 ± 0.96 hours) **(Fig.1B)**.

We additionally evaluated the variance between researchers for each brain area and mouse. For this evaluation, images with poor background fluorescence were excluded from the analysis. We found that AxoDen had lower inter-researcher variability compared to Mean Intensity analysis (AxoDen: 5.42 ± 0.58; Mean Intensity: 16.01 ± 3.69; p-val = 0.05) **(Fig. lC)**. Evaluation of the variance for each brain ROI clearly shows that variance between researchers was generally lower for AxoDen **(Fig. 1D)** compared to Mean Intensity analysis **(Fig. lE)**. These results are further consolidated when comparing the data collected by each experimenter. The values obtained were more similar between experimenters when using AxoDen **(Fig. lF)** than when using Mean Intensity analysis **(Fig. lG)**. Altogether, this comparison shows that 1) Axoden notably decreases the time required to collect and analyze the data and 2) that Axoden reduces inter-experimenter variability.

**Figure 1.**
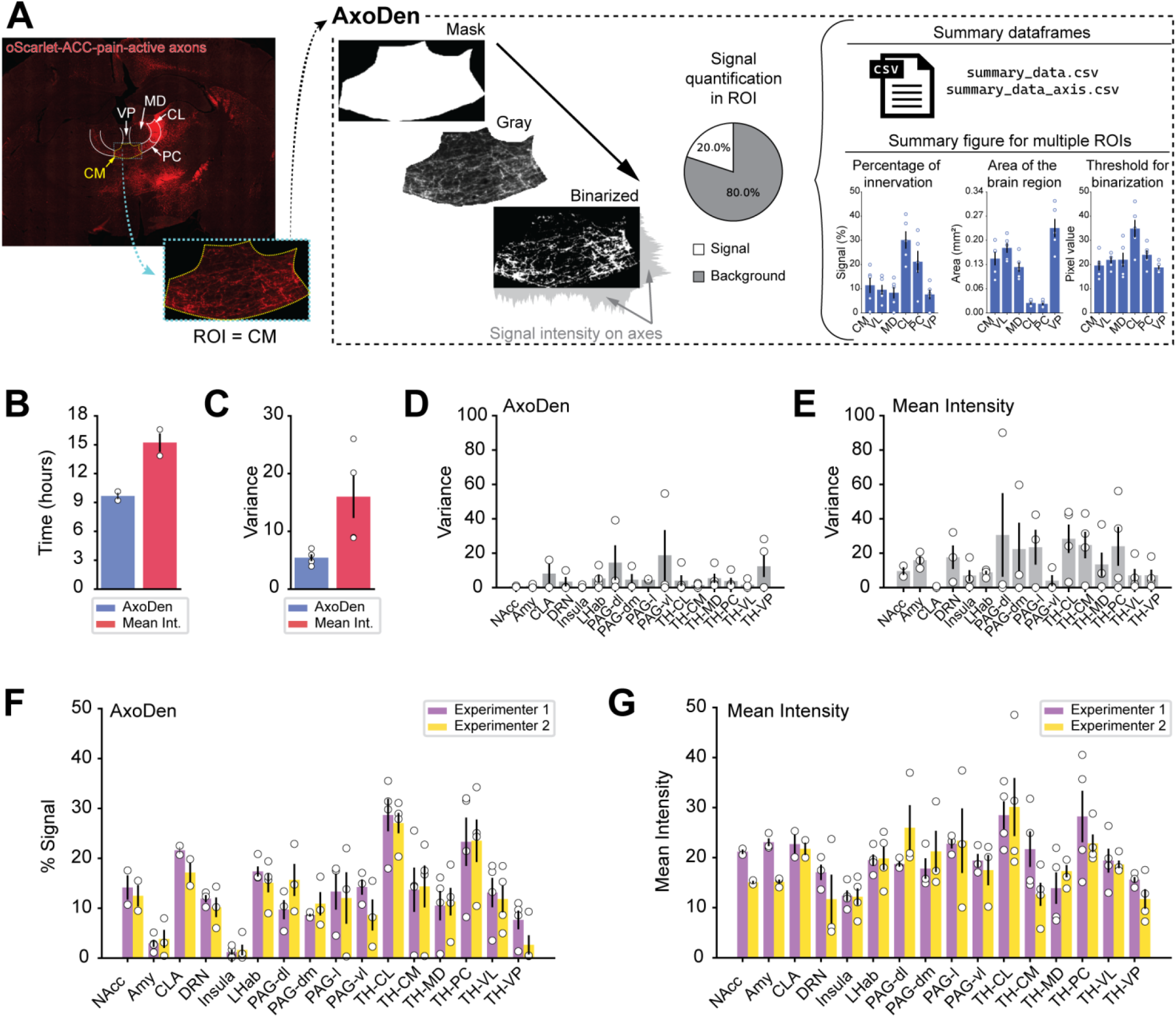
AxoDen overview and comparison to Mean Intensity method. **A)** Overview of AxoDen steps and outcome using example oScarlet-ACC-pain-active axons projecting to midline thalamic nuclei. **B)** Time needed to collect the data for AxoDen and Mean Intensity. **C)** Overall variance between researchers for AxoDen and Mean Intensity. **D)** Variance between experimenters when using AxoDen. **E)** Variance between experimenters when using Mean Intensity. F) Percent of signal in different ROIs indicating the percent of the ROI receiving axonal projections for each experimenter using AxoDen. **G)** Mean fluorescence Intensity values collected for each brain area by different experimenters. B) Each dot represents one experimenter. **A, C-G)** Each dot represents one animal. All values are mean ± SEM. ACC, anterior cingulate cortex; Amy, amygdala; CL, centrolateral nucleus; CLA, claustrum; CM, centromedial nucleus; dl, dorsolateral; dm, dorsomedial; DRN, dorsal raphe nucleus; I, lateral; LHab, lateral habenula; MD, mediodorsal nucleus; NAcc, nucleus accumbens; PAG, periaqueductal gray; PC, paracentral nucleus; PV, paraventricular nucleus; ROI, region of interest; TH, thalamus; vl, ventrolateral; VL, ventrolateral nucleus; VP, ventroposterior nucleus.

### Experiment Overview for Axonal Quantification using AxoDen

AxoDen has been designed to provide a streamlined analysis of axonal projections in anatomical studies. Therefore, the work-flow for quantifying axonal innervations of pre-defined brain ROIs follows a four-step process **(Fig. 2):**

**Figure 2.**
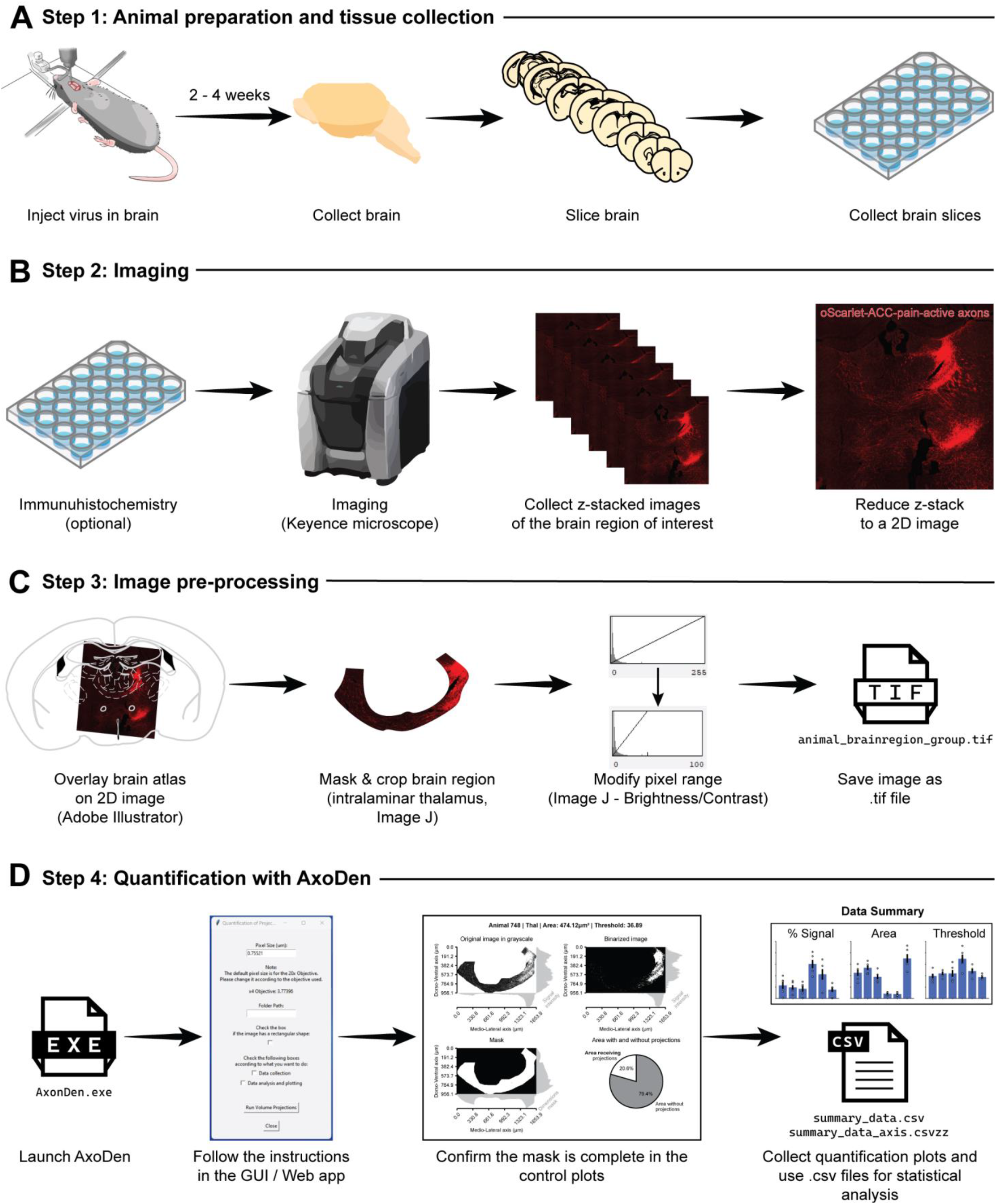
Experimental workflow for anatomical analysis of axonal projections in the intralaminar thalamus. **A)** Steps for animal preparation and tissue collection. **B)** Steps for image acquisition. **C)** Steps for image pre-processing.**D)** Steps for axonal quantification with AxoDen and its outputs.

## Step 1: Animal preparation and tissue collection (Fig. 2A)

### 1. Viral Vector Injection

Administer a viral vector containing a fluorescent reporter into the targeted brain region of the animal model of choice to label axonal projections.

### 2. Expression Period

Allow a∼4-week period for optimal viral expression in neuronal projections.

### 3. Euthanasia and Brain Harvesting

Euthanize the animal following approved ethical guidelines and carefully extract the brain.

### 4. Tissue Sectioning

Slice the brain into sections ranging from 30 to 50 pm in thickness, suitable for subsequent imaging.

## Step 2: Image acquisition (Fig. 2B)

### 1. Immunohistochemistry (Optional)

If necessary, apply immunohistochemistry techniques to amplify the fluorescent signal to ensure visibility - or increase the signal-to-noise ratio - of axonal projections.

### 2. Z-Stack Imaging

Capture the axonal projections at multiple depths of the brain ROI by acquiring z-stacks using a 20x objective lens on a microscope.

### 3. Image Processing

Process the z-stacks using a maximal projection mode to generate 2D images that minimize background fluorescence while preserving signal integrity.

## Step 3: Image pre-processing **(Fig. 2C)**

### 1. Atlas Overlay

Superimpose the relevant brain atlas onto your images to accurately identify the ROls for the later masking step.

### 2. Region Masking

Precisely delineate, mask, and crop the brain area of interest, guided by the overlay created previously to ensure accurate measurement zones. We-recommend this step to be performed in software such as lmageJ.

### 3. Fluorescence Intensity Adjustment (Optional)

Adjust the fluorescence intensity to increase the signal-to-noise ratio without completely eliminating background fluorescence, ensuring that:

a. All axonal projections are brighter than the background.
b. The background does not contain black pixels.

For details read the sections Consideration 1 and 2.

### 4. Image Rotation (Optional)

Rotate the image to align the media-dorsal axis to the x axis and the dorso-ventral axis to they axis. This step can allow for a detailed study of differential innervation of cortical layers. For details read the section Consideration 3.

### 5. Image Batch Naming

*Save* all processed and cropped images in the format animallD_brainregion_group_[additional_information].tif to follow the data management organization of the algorithm for batch processing. For details read the section Consideration 4.

## Step 4: Quantification with AxoDen **(Fig. 2D)**

### 1. Script Initialization

Launch AxoDen in any of its available forms.

### 2. Provide the requested information to the GUI

a. Pixel Size: The default pixel size is for a 2ox objective [Nikon PlanApo 2ox 0.75NA/0.60mm working distance] in a Keyence microscope [BZ-X Series].
b. Input the data:
  i. Users of the web app, upload the images to analyze.
  ii. Users of the executable provide the directory path containing the saved images to the script interface.
  iii. Users of the Python package or the cloned repository can use either of the methods to input their data.
c. Toggle option if the images are masked and cropped to a defined region of interest.

### 3. Analysis Execution

*Run* the scriptto initiate automated axonal quantification. In the stand-alone GUI, the control plots, summary figures and .CSV files containing the dataframes generated during the analysis will be saved in the provided output folder. In the Web application, the user can decide which information to download.

### 4. Check the control plots

Make sure the mask covers the entire area of the brain ROI.

### 5 Collect the data of interest from the output files

a. CSV files contain the raw data used to create the figures and can be used by the user for later statistical anal-sis:
  i. One CSV file provides information on the overall innervation of each brain area and animal.
  ii. The second CSV file provides information on the fluorescence intensity along the X and the Y axis of the images.
b. PDF files contain figures that the user can open and modify in Adobe Illustrator:
  i. The user can use the control plots for each animal and brain regions to confirm each file has been adequately processed.
  ii. The summary plot provides information on the overall innervation density, area (in mm^2^) and threshold used for binarization for each brain region and informs of the number and identity of the animals included in the analysis.

### AxoDen algorithm workflow

The AxoDen algorithm processes images that have been previously masked and cropped to specific shapes, as well as unmasked images. Masked and cropped images are conventionally treated as rectangular frames, with areas lacking tissue represented by black pixels or a value of zero **(Fig. 3A)**. This format is used to retrieve the mask, which delineates the boundaries of the chosen brain region, thereby differentiating between informative (tissue-containing) and non-informative (masked-out) segments of the image **(Fig. 3B)**.

**Figure 3.**
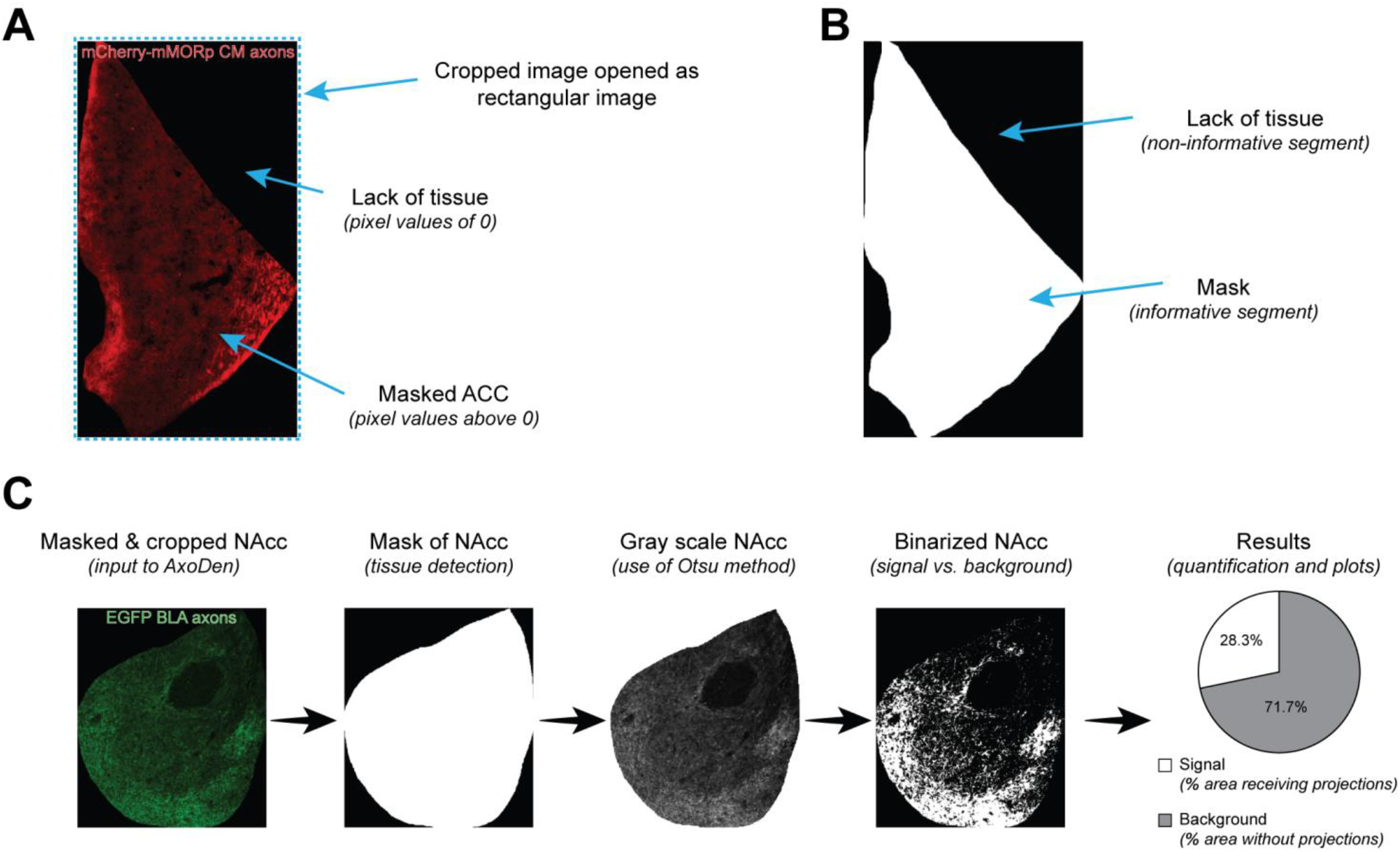
AxoDen workflow with masked and cropped images. **A)** Components of a masked and cropped image of the ACC receiving CM axons positive for mu opioid receptors (MOR) labeled with mCherry. **B)** Illustration of the detected mask by AxoDen from the masked and cropped ACC image. **C)** AxoDen workflow for an image of the masked and cropped nucleus accumbens (NAcc) with projections labeled with GFP.

This analysis is independent of the fluorophore used to label projections, given that AxoDen converts all input images into gray scale **(Fig. 3C)**. After images have been transformed to grayscale, the Otsu method (Otsu, 1979) is applied using the filters.threshold_otsu function from the scikit-image Python library, which calculates an optimal threshold by finding a value that maximizes the variance between classes. In our case, classes are “signal” and “background”. The resulting threshold is applied to binarize the gray scale image such that pixels above the threshold are classified as signal and given a value of 1, and pixels below the threshold are considered background and are assigned a value of o, ensuring signal is segregated from background. Taking the binarized image, AxoDen calculates the percentage of the brain region identified with the mask that corresponds to signal versus background, thereby offering an overall view of the percent of innervation of the ROI **(Fig. 3C)**.

AxoDen additionally offers the possibility to collect the signal intensity levels longitudinally (x-axis) and latitudinally (y-axis), offering insights into the innervation across different anatomical axes when the brain region is appropriately aligned with the media-lateral and ventro-dorsal axes. Read Consideration 3 for this optional feature.

### Availability

Scientific research has long faced a reproducibility crisis, primarily due to the challenges in replicating experimental conditions and analyses across different laboratories. To address this issue, it is crucial to develop standardized procedures and analyses that are user-friendly for the entire scientific community. AxoDen has been designed with enhanced usability and accessibility in mind, standardizing the quantification of axonal projections. As a result, AxoDen is freely available to all scientists, regardless of their familiarity with analysis scripts or coding languages.

AxoDen can be used in three distinct modes, ranked here from requiring the least coding expertise to the developer level, for those who need to modify or integrate AxoDen’s functions into their own scripts:

### 1. Web Application

AxoDen is accessible as a web application to anyone through a web browser without signup or user account required. This offers a platform-independent solution that performs the computations in the cloud, broadening AxoDen usability. Any image uploaded here is not saved in any web server, and remains in memory only for the analysis, ensuring the confidentiality of any sort of data the user analyzes. The web application can be found here: https://axoden.streamlit.app/.

### 2. Executable

AxoDen is available as a stand-alone graphic user interface (GUI) for Windows, MacOS and Ubuntu/Linux. These stand-alone AxoDen GUI versions can be downloaded here: https://github.com/raqueladaia/AxoDen/releases as zip files. After extracting the content of the zip files, the executable will be ready to launch by double clicking on it. The messages that appear in the stand-alone GUI inform about the steps the algorithm is performing.

### 3. Source Code

For those researchers who wish to inspect the source code, the underlying scripts of AxoDen can be accessed by either 1) Cloning of GitHub Repository (https://github.com/raqueladaia/AxoDen), or by 2) installing the Python pip package axoden (https://pypi.org/project/axodenD This flexibility allows for tailored modifications of the algorithm to suit specific research needs.

Each function of AxoDen is documented in the documentation website of AxoDen: https://raqueladaia.github.io/AxoDen/. The instructions on how to use, install or download AxoDen can be found in the GitHub repository. In the Web App application, a “How To” section provides instructions on how to prepare for and use AxoDen on the web browser. Using AxoDen via cloning of the repository or the pip installation package gives access to both the GUI and the local use of the web application. These diverse formats enhance AxoDen’s accessibility and adaptability, catering to varying user preferences and technical infrastructures.

### Considerations for proper use of AxoDen

Every digital image is a mosaic of pixels, each encoding information about color and intensity. Commonly, cameras produce color images by synthesizing three primary colors– referredto as channels: Red, Green, and Blue (RGB). Hence, an image inherently possesses three dimensions: 1) width (x-axis), 2) height (y-axis), and 3) the color channels (z-axis). This structure allows us to conceptualize an image as a three-dimensional matrix, wherein each pixel is represented by a trio of values that correspond to the intensity levels of Red, Green, and Blue. The linear combination of the intensity values of the various channels give rise to the wide spectrum of colors we observe in an image. Typically, images are captured in an 8-bit format, meaning the pixel intensity in each channel can assume 28, or 256, possible values ranging from o to 255 in steps of 1. In this format, a value of o across all channels results in black, while a value of 255 across all channels manifests as white.

With these fundamentals in place, adhesion to the first two **considerations ensures precise axonal detection**.

### Consideration 1: Avoid overexposure during image acquisition

During image acquisition under a microscope, it is common practice to adjust the brightness to enhance the visibility of the fluorophore. Increasing the ‘Brightness’ - or ‘Exposure’ - parameter extends the duration for which the sample is illuminated by the excitation light-this is known as exposure time-, thereby improving signal-to-noise ratio. Nevertheless, excessive exposure causes two significant issues: photobleaching and image oversaturation. Photobleaching is a reduction in fluorophore emission intensity due to an irreversible photochemical reaction following extended exposure to light. Image oversaturation, or overexposure, occurs when too many pixels reach the maximum intensity value of 255, rendering those regions devoid of usable data, and leading to an overestimation of signal **(Fig. 4 A, B)**. Consequently, it is crucial to limit the number of pixels reaching their maximum value.

**Figure 4.**
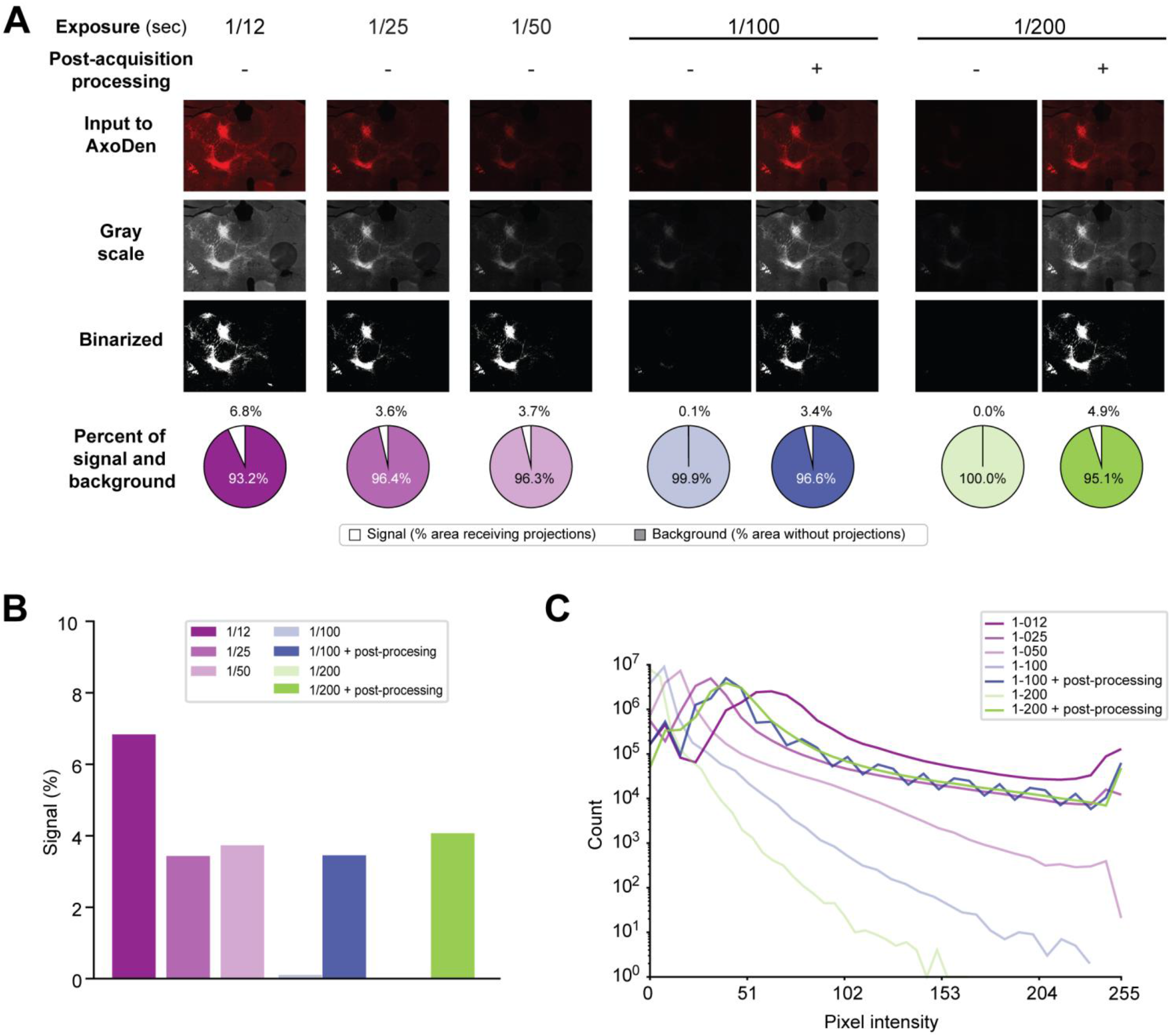
The effect of exposure time during acquisition. **A)** Series of images of a-Scarlet-labeled projections from the ACC to the thalamus acquired at different exposure times, with (+)or without(-) post-acquisition image processing, and their transformations and quantification of signal by the AxoDen algorithm. **B)** Comparison of the percent of innervation measured for each acquisition time. **C)** Histograms of the pixel intensity values for each acquired image.

Thanks to the dynamic thresholding step in AxoDen, different exposure times do not affect signal detection, as seen in the example exposures times of 1/25 and 1/50 seconds **(Fig. 4A, B)**. Lower exposure times, such as 1/100 and 11200 seconds, do not provide enough contrast for AxoDen to segregate signal from background. However, since the pixel information remains in the image, post-acquisition enhancement of brightness can achieve the same quantification values at those acquired at 1/25 and 1/50 seconds **(Fig. 4B)**. This is evident when exploring the histogram of images acquired at different exposure times **(Fig. 4C)**. Decreasing the exposure times shifts the distribution of pixel intensity values towards the left, increasing the number of pixels with low values. Increasing the brightness post-acquisition shifts the distribution of pixels towards the right, increasing the number of pixels with high values and allowing AxoDen to distinguish between “signal” and “background” fluorescence. Therefore, we recommend keeping exposure times at low levels, especially when intensity levels can be enhanced post-acquisition **(Fig. 5)**.

**Figure 5.**
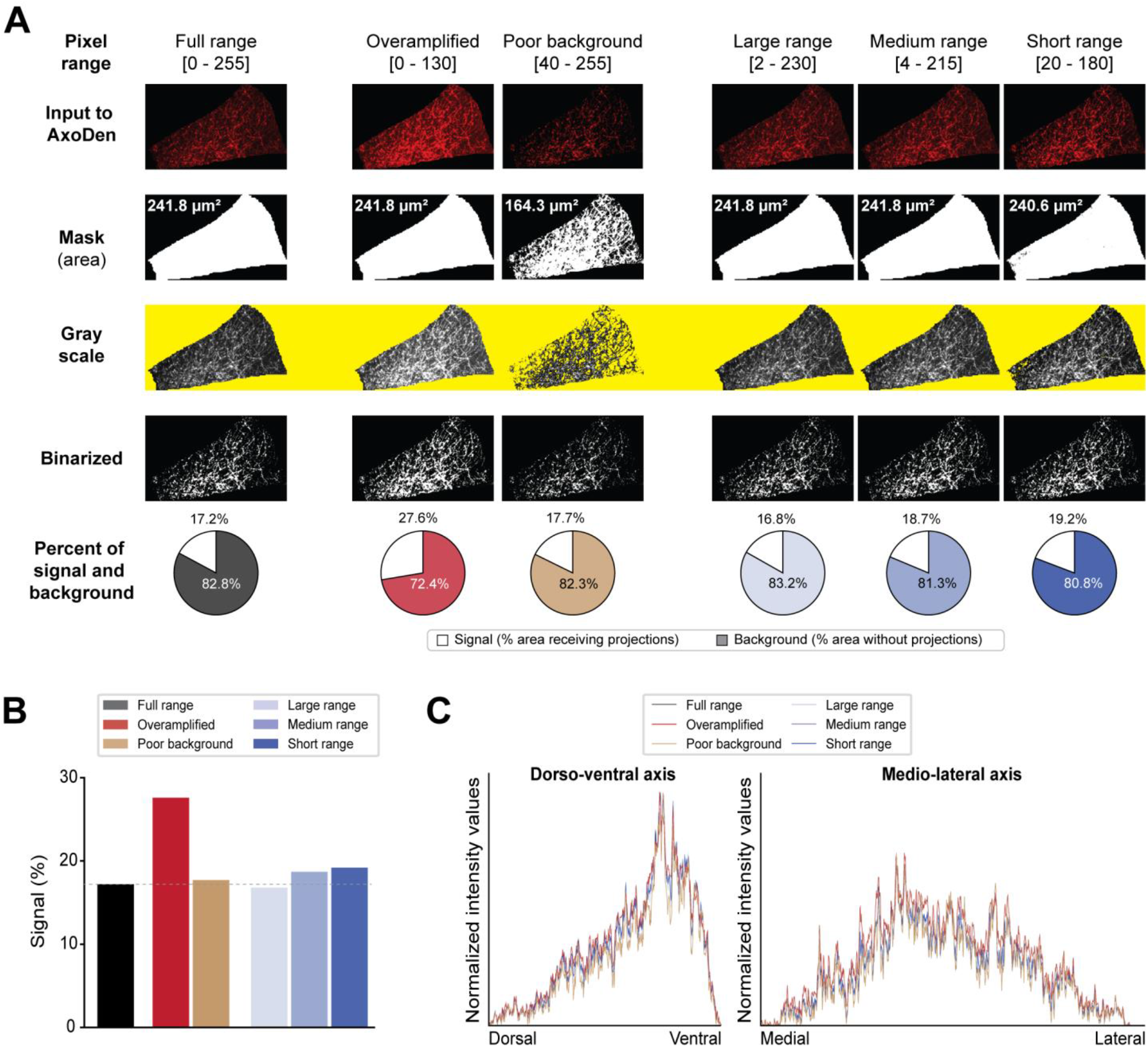
Effect of pixel ranges on the signal detection capabilities of AxoDen. **A)** Series of images of oscarlet-labeled projections from ACC to PAG-vl with different pixel ranges and their transformations by the AxoDen algorithm. The yellow band on the gray scale row is used to visualize the transparency of the gray scale images caused by an incomplete mask. **B)** Overall percentage of signal for each condition. The dotted line is used as a reference for the signal% computed for the full range image. **C)** Normalized intensity for dorso-ventral and media-lateral axes.

### Consideration 2: Preserve background fluorescence in the sample area

In scenarios where the analysis algorithm processes masked images, maintaining a detectable level of background fluorescence– denoted by pixel values greater than zero– is crucial. Due to the convention of storing images in rectangular formats, areas beyond the periphery of the masked sample are assigned a pixel value of zero, indicative of an absence of data **(Fig. 3A)**. Consequently, our algorithm discerns zero values as void of informational content **(Fig. 3B)**. Fig. 5 shows the effect of different postacquisition adjustments of an image where the ventrolateral region of the periaqueductal gray (PAG-vl) has been masked and cropped. Keeping the full range of pixels **(Fig. SA, column 1)**, AxoDen correctly classifies the “informative” pixels from the “non-informative” pixels creating a mask that covers the full area of PAG-vl **(Fig. SA, column 1, second row)**. Obtaining a mask that covers the full brain region **(Fig. SA, column 1**,**third row)** confirms that AxoDen exclusively considers the pixels that fall within the mask **(Fig. 3A)** in order to transform the image into gray scale **(Fig. SA, column 1, third row)** and compute the dynamic threshold to transform the image into a binary image **(Fig. SA, column 1, fourth row)**. The binary image will serve the user to confirm that there is no bleeding of the signal and to quantify the percent of pixels that have been identified as “signal” and “background” **(Fig. SA, column 1, fifth row)**.

Often times, images may present low pixel intensity levels because the exposure time during acquisition under the microscope has been kept low to preserve the fluorophore integrity. Therefore, a commonly used post-acquisition strategy consists in increasing the signal-to-noise ratio (SNR) by adjusting the Brightness/Contrast settings. Such modification decreases the image’s dynamic range by clipping the ends of the pixels value spectrum. In other words, the spectrum of potential pixel intensity values is narrowed **(Fig. SA, columns 2 to 5)** and therefore any value below the new minimum is interpreted as o, and any value above the new maximum is interpreted as 255. However, care must be taken to prevent the overamplification of the signal by clipping the maximum values **(Fig. SA, column 2)**. In the example provided, the new maximum is set to 130, and therefore any pixel with a value above 130 is considered as 255. This extreme modification, which we name overamplification, leads to the artificial spread of signal intensity to adjacent pixels, falsely inflating the perceived axonal density **(Fig. SA, column 2, last row)**. Another crucial aspect that ensures the algorithm’s accurate interpretation and analysis is the retention of minimal background fluorescence within the sample **(Fig. SA, column 3)**, meaning that the pixel values within the ROI remain above zero. This prevents the erroneous classification of sample as absence of data. Increasing the minimum of the pixel range from o to 40 sets all pixels with values below 40 as the new o. Therefore, all pixels that have acquired a new value of zero are considered as “non informative” **(see Fig. 3B)** and are left out of the analysis, resulting in a mask of a smaller area compared to the area of the brain ROI **(Fig. SA, column 3, second row)**. This in turn generates a “patchy” gray-scale image of a smaller area - appreciated by the yellow background appearing throughout the vlPAG - **(Fig. SA, column 3, third row)**, ultimately yielding misleading quantitative results. Nonetheless, clipping the full pixel range at different minimum and maximum levels yielding images with new pixel ranges of different lengths **(Fig. SA, columns 4 to last)** shows that the algorithm is robust to variations of pixel ranges **(Fig. SA, columns 4 to last)**. However, when the clipping is so severe that the background fluorescence reaches values of zero and the mask does not cover the entirety of the brain region, an overestimation of the signal occurs **(Fig. SA, last column)**. Comparison of the signal percentage in the previous conditions shows that images with overamplified signal yield the highest overestimation of projections **(Fig. SB)**. However, variation of the signal percent is kept low between the different pixel ranges and the normalized intensity across both the dorso-ventral and the media-lateral axes do not show large differences between conditions **(Fig. SC)**. The robustness of AxoDen to acquisition settings **(Fig. 4)** as well as to post-acquisition adjustments **(Fig. 5)**, which is provided by the dynamic thresholding method, prevents inter-researcher variation **(Fig. 1)**, making AxoDen a tool easily accessible to researchers at all career levels.

### Consideration 3 (Optional): Rotate the image for accurate spatial distribution of axes

In contexts requiring detailed spatial characterization,such as the evaluation of innervation profiles across cortical layers, the user can opt to rotate the image, previous to masking and cropping the region of interest, and feed the resulting image to AxoDen **(Fig. 6)**. This alignment ensures that the fluorescence quantification accurately reflects the spatial distribution of interest, thereby yielding data that are both reliable and biologically meaningful. With the example of projections from the basolateral amygdala (BLA) to ACC (Wojick et al., 2024), the user rotates the image to align the dorso-ventral and media-dorsal axes with the y and x axes, respectively. Then, proceeds to mask and crop the ACC **(Fig. SA)** to later feed the resulting image into AxoDen. AxoDen transforms the image into gray scale and later into a binary image, calculating the signal intensity along the x and y axis for each transformation, as well as the percentage of total signal detected in the binary image **(Fig. 6B)**. The intensity profiles for the gray scale and the binarized image are different, showing larger variation in that of the binarized image **(Fig. SC)**. Overlay of the intensity profiles of the media-lateral axis with the cropped ACC shows that the profile obtained from the binarized image better recapitulates the innervation pattern of the different cortical layers of ACC **(Fig. 6D)**. Therefore, only the information of the binarized image should be taken into consideration for this analysis, given that in the background fluorescence of gray-scale images occludes the innervation patterns of different cortical layers.

**Figure 6.**
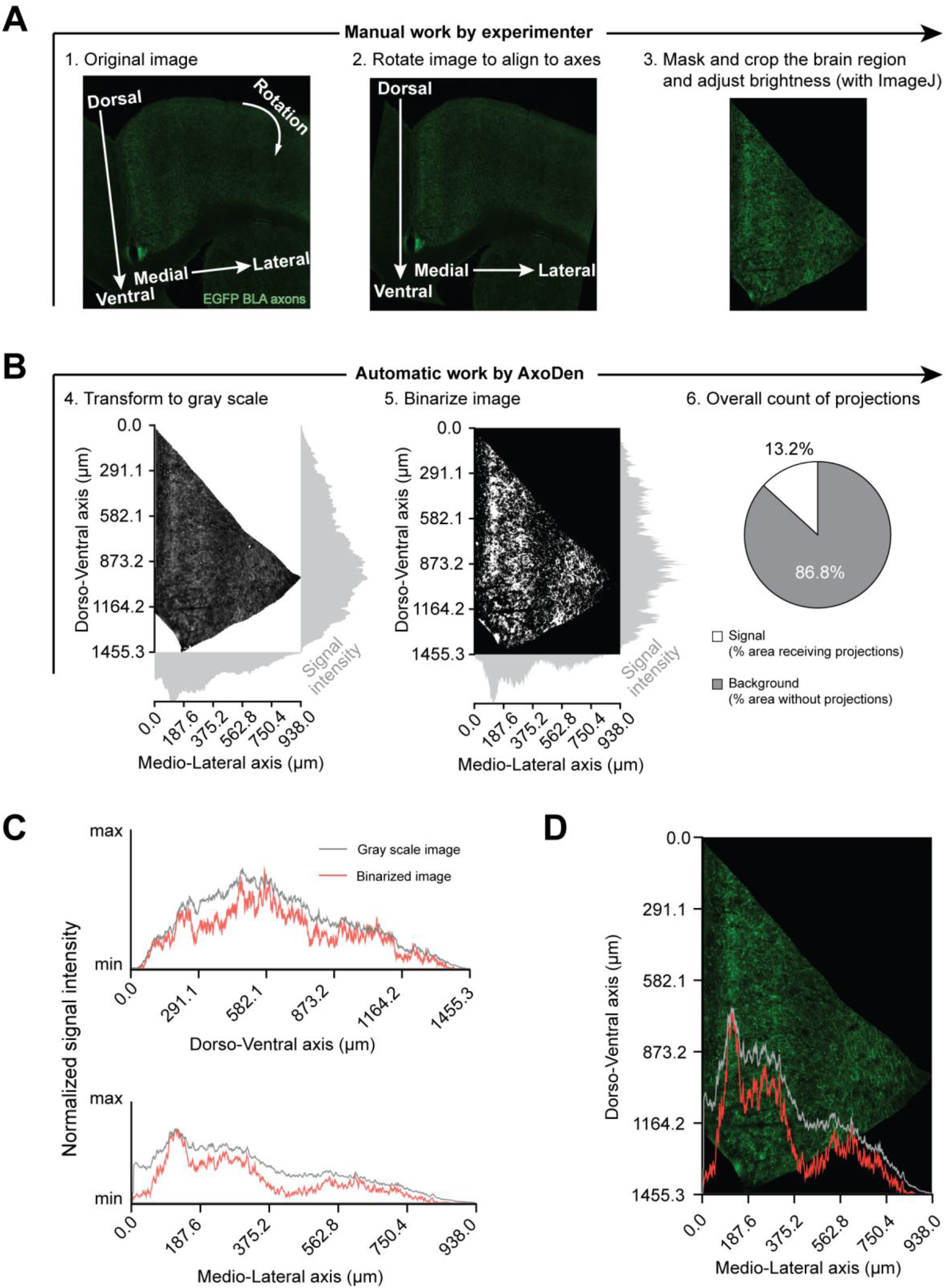
Spatial distribution of projections along the x and y axes in the ACC. **A)** Manual steps users need to follow to first rotate and then crop an image to align the dorso-ventral and the media-lateral axes to they and x axis, respectively. **B)** Automatic AxoDen steps where the signal intensity is measured along the axes and globally. **C)** Normalized signal intensity along the dorso-ventral and the media-lateral axes. D) Overlay of the signal intensity on the media-lateral axis with the cropped image.

### Consideration 4: Adhere to the algorithm’s naming conven tion

Any algorithm operates through a sequence of systematically arranged instructions. Thus, adhering to the AxoDen’s naming convention is crucial to guarantee that data from each image are accurately extracted and categorized for effective analysis. AxoDen relies on the file name to derive essential details about each image, using the underscore symbol (“_”) as a delimiter to segregate distinct information details within the name. Subsequently, it categorizes the segmented information based on the position it occupies. The first position is designated as the subject identifier, the second position denotes the brain region, and the third position represents the experimental group. In cases where there is only one experimental group, write the same word (i.e. “group”) in the third position of the file name for all the images of the given experiment. For instance, in a filename like:

~~~
mouse123_AnteriorCingulateCortex_control_addi-
tional-information.tif
~~~

`mouse123` is interpreted as the subject identifier, `AnteriorC-ingulateCortex` specifies the brain region and `control` identifies the experimental group. The algorithm disregards any text succeeding the experimental group identifier, providing a series of positions for supplementary details about the image that does not influence the core data extraction process. Furthermore, the algorithm recognizes and processes images specifically in the `.tif` format. Consequently, ensuring that all images prepared for analysis are saved in this format is imperative.

## Discussion

Here we introduce AxoDen **(Fig. lA)**, a platform to streamline, standardize and speed up the quantification of axonal innervation of defined brain ROls. AxoDen is robust against different acquisition times **(Fig. 4)** and diverse post-acquisition adjustments **(Fig. 5)**. This robustness decreases the variability in the measurements taken by different researchers **(Fig. 1 C-G)**. AxoDen additionally provides a more accurate representation of brain area innervation as it quantifies the distribution of axonal projections in the x and they axes **(Fig. 6)**. Furthermore, given that this method relies on the range of pixel values of one single channel, the color of the fluorophore does not impact the quality of the quantification **(Fig. 3)**. Thus, our method promises to make high quality analysis of axonal innervation accessible to every laboratory and researcher irrespective of training experience, and to enhance the precision of axonal density quantification for any image of fluorescently labeled axonal projections taken with an objective of 2ox or above for any animal species.

### Expertise required to implement AxoDen

AxoDen is versatile, capable of analyzing any brain region independently of its shape, fluorophore used or animal species. It is specifically designed to accommodate users with varying levels of coding expertise, thereby broadening its applicability across diverse research contexts. As such, users are not required to have advanced data analysis skills to utilize the software effectively. However, a fundamental understanding of neuroanatomy for the specific species under investigation is essential. This knowledge is crucial to 1) accurately identify and select the targeted brain region for analysis, and 2) correctly orient the image to align with the relevant axes, ensuring that the derived x-and y-axis fluorescence intensity profiles are biologically significant. Additionally, users should be familiar with basic image analysis software, such as the widely used lmageJ, to adjustthe signal-to-noise ratio. Enhancing this ratio has the potential to improve the clarity and distinguishability of the signal, facilitating more accurate and reliable quantification of axonal density, for those images with low exposure times.

### Limitations

AxoDen provides notable improvements in the quantification of axonal projections, yet it has limitations that need attention. First, while AxoDen effectively quantifies axonal projections, it cannot distinguish axons from cell bodies. As a result, neuronal somas might be erroneously included in the axonal counts, potentially biasing the outcome. As with somas, AxoDen is unable to identify and exclude artifacts like bubbles or dust. To address these challenges, a recommended preprocessing step involves manual identification and removal of neuronal cell bodies and artifacts from the images before proceeding with the analysis. Although this step introduces additional time investment, it substantially enhances the precision of the axonal quantification. Another limitation arises from the method’s optimization for images obtained with a 2ox objective lens, which strikes a balance between field of view and resolution sufficient for identifying axonal shapes and assessing their density and distribution. However, this specificity restricts the method’s applicability to images captured at this magnification. Utilizing images obtained with a 4x objective lens is not advised due to the lower resolution, which impedes the accurate differentiation of axons from other structures, thereby compromising the integrity of the analysis. For studies necessitating the use of 4x magnification– often aimed at broader regional assessments– we suggest adhering to traditional mean fluorescence intensity measurements, despite their known limitations. This recommendation is made with the understanding that the insights gained from such analyses at lower magnifications serve different research objectives, primarily related to general patterns of innervation rather than detailed quantifications of axonal density and distribution.

## Methods

### Animals

All experimental procedures were approved by the Institutional Animal Care and Use Committee of the University of Pennsylvania and performed in accordance with the US National Institutes of Health (NIH) guidelines. Mice aged 2-5 months were housed 2-5 per cage and maintained on a 12-hour reverse light-dark cycle in a temperature and humidity-controlled environment with adlibitum food and water.

Mice genetic background was the following for each set of images:

- Red-fluorescently-labeled axons of the images used for the validation of AxoDen were from TRAP2 mice crossed with CAG Sun1 (B6;129-Gt(ROSA)26Sortm5(CAG-Sun1/sfGFP)Nat/J) reporter mice that express a GFP fluorophore in a ere-dependent manner (“TRAP2:CAG-Sunl sfGFP”). These were purchased from Jackson Laboratory, strain *ft* 021039 and bred to homozygosity for both genes.
- Images of green-fluorescently-labeled axons were from TRAP2 mice crossed with Ai9 (B6.Cg-Gt(ROSA)26Sortm9(CAG-tdTomato)Hze/J) reporter mice expressing a tdTomato fluoro-phore in a ere-dependent manner (“TRAP2:tdTomato”). Purchased from Jackson Laboratory, strain *ft* 007909 and bred to homozygosity for both genes.
- C57BL/6J wild type mice were used for the image of axons innervating the ACC and were purchased from Jackson Laboratory, strain *ft* 000664.

### Viral vectors

All viral vectors were either purchased from Addgene.org, or custom designed and packaged by the authors as indicated. All AAVs were aliquoted and stored at -so_0_c until use and then stored at 4°C for a maximum of four days.

The next two AAVs were used for the red-fluorescently-labeled axons of the images used for the validation of AxoDen:

- AAV5-mMORp-FlpO (Deisseroth lab, Stanford; titer: 1.9e12 vg/mL)
- AAV8-Ef1a-Con/Fon-oScarlet (Addgene 137136-AAVS; titer: 2.2e12 vg/mL)
- The following AAV was used for the green-fluorescently-labeled axons:
- AAV5-hSyn-DIO-EGFP (Addgene 50457-AAV5; titer: 1.3e12 vg/mL)
- For the image of axons innervating the ACC, the following AAVs were used:
- AAV1-mMORp-hM4Di-mCherry (Deisseroth lab, Stanford; titer: 1.17e12 vg/mL)

### Stereotaxic surgery

Adult mice (∼8-10 weeks of age) were anesthetized with isoflurane gas in oxygen (initial dose = 5%, maintenance dose = 1.5%), and fitted into Kopf stereotaxic frames for all surgical procedures. 10 µL Nanofil Hamilton syringes (WPI) with 33 G beveled needles were used to intracranially infuse AAVs into the different brain areas of interest. Based on the Paxinos mouse brain atlas, the following coordinates were used for each set of images:

- Images of red-fluorescently-labeled axons used for the validation of AxoDen:ACC: AP: -1.50 mm, ML:± 0.3 mm, DV: -1.5 mm
- Images of green-fluorescently-labeled axons: BLA: AP: -1.20 mm, ML: 3.20 mm, DV: -5.20 mm
- Image of axons innervating the ACC, the following AAVs were used: CM: AP: -1.70 mm, ML: 0.70 mm, DV: -4.00 mm, angle: 10°.

Mice were given a 3-8-week recovery period to allow ample time for viral diffusion and transduction to occur. For all surgical procedures, meloxicam (5 mg/kg) was administered subcutaneously at the start of the surgery, and a single 0.25 ml injection of sterile saline was provided upon completion. All mice were monitored and given meloxicam for up to three days following surgical procedures.

### TRAP protocol (tamoxifen induction)

The images of red-fluorescently-labeled axons used for the validation of AxoDen were from pain-active neurons in the anterior cingulate cortex (ACC) (James et al., 2024), labeled via tamoxifen induction. Mice were habituated to the testing room the day before TRAP execution and no nociceptive stimuli were delivered. On both days of habituation and TRAP procedure, mice were placed within red plastic cylinders (10.16-cm in diameter), with a red lid, in a raised metal grid. On the day of the TRAP procedure, mice were habituated to the room and the cylinder for 60 minutes and then they received 20 stimuli consisting of a water drop at 55 °c interspaced by 30 seconds over 10 minutes. Following the stimulation, the mice remained in the cylinder for an additional 60 min before injection of 4-hydroxytamoxifen (20 mg/kg in ∼ 0.25-mL vehicle; subcutaneous). After the injection, mice remained in the cylinder for an additional 2 hours to match the temporal profile for c-FOS expression, at which time the mice were returned to the home cage.

### Tissue processing

Animals were anesthetized using FatalPlus (Vortech) and transcardially perfused with 0.1M phosphate buffered saline (PBS), followed by 10% normal buffered formalin solution (NBF, Sigma, HT501128). Brains were quickly removed and post-fixed in 10% NBF for 24 hours at 4 °c, and then cryo-protected in a 30% sucrose solution made in 0.1 M PBS until sinking to the bottom of their storage tube (∼48 h). Brains were then frozen in Tissue Tek O.C.T. compound (Thermo Scientific), coronally sectioned on a cryostat (CM3050S, Leica Biosystems) at 30 µm and the sections stored in 0.1 M PBS.

### Fluorophore amplification

Floating sections were permeabilized in a solution of 0.1 M PBS containing 0.3% Triton X-100 (PBS-T) for 30 min at room temperature and then blocked in a solution of 0.3% PBS-T and 5% normal donkey serum (NDS) for 2 hours before being incubated with primary antibodies in a 0.3% PBS-T, 5% NDS solution for ∼16 h at room temperature. Following washing three times for 10 min in PBS-T, secondary antibodies prepared in a 0.3% PBS-T, 5% NDS solution were applied for ∼2h at room temperature, after which the sections were washed again three times for 5 mins in PBS-T, then again three times for 10 min in PBS-T, and then counterstained in a solution of 0.1 M PBS containing DAPI (1:10,000, Sigma, D9542). Fully stained sections were mounted onto Superfrost Plus microscope slides (Fisher Scientific) and allowed to dry and adhere to the slides before being mounted with Fluoromount-G Mounting Medium (lnvitrogen, 00-4958-02) and cover slipped.

For each batch of images, the antibodies included:

- Images of red-fluorescently-labeled axons of the images used for the validation of AxoDen:
  - 1°Ab: rabbit anti-dsRed [1:1000, Takara, 632496]
  - 2°Ab: Alexa-Fluor 555 donkey anti-rabbit [1:500, lnvitrogen Thermo Fisher, A31572]
- Images of green-fluorescently-labeled axons:
  - 1°Ab: chicken anti-GFP [1:1000, Abeam, ab13970]
  - °Ab: Alexa-Fluor 488 donkey anti-chicken [1:500, Jackson lmmuno, 703-545-155]
- Image of red-fluorescently-labeled axons innervating the ACC:
  - 1°Ab: rabbit anti-dsRed [1:1000, Takara, 632496]
  - 2°Ab: Alexa-Fluor 594 donkey anti-rabbit [1:500, Abeam, A21207].

### Imaging

All images were acquired with the 2ox objective Nikon PlanApo 2ox 0.75NA/0.60mm working distance, using the Keyence microscope BZ 8lOX.

### Datasets

All images were generated in the Corder Lab as described in the methods above.

### Axonal quantification using mean fluorescence intensity

The acquired images were opened in FIJI. The brightness of the image was increased to better visualize the fluorescence of the axons. The ROI selection tool shaped as a rectangle was selected. A rectangle was placed over the brain region of interest so that the area of the rectangle occupied most of the area of the brain region, without exiting the boundaries of the brain region. The brightness was restored to its original value, and it was adjusted to increase the signal-to-noise ratio within the rectangle. The “Histogram” function within the menu command “Analyze” was used to collect the mean fluorescence intensity within the rectangle by collecting the value “Mean”. Data on the mean intensity fluorescence were stored in Excel files.

## Acknowledgements

This work was funded by NIH NIGMS DP2GM140923, R01DA056599, R01DA054374, R01NS130044 (G.C.). We thank the University Laboratory Animal Resources (ULAR) group at the University of Pennsylvania for assistance with rodent husbandry and veterinary support. The authors express their gratitude to Corinna Oswell for her contribution of images to the dataset utilized for validation, to Jessica Wojcik for supplying axon images labeled with green fluorescent protein, and to Lindsay Ejoh for providing the anterior cingulate cortex image labeled with mCherry.

## Author contributions

R.A.S.O conceptualized and planned out the algorithm for AxoDen, performed data collection, analysis and visualization, supervision, wrote the original manuscript and edited and revised the manuscript. E.L and O.J validated AxoDen, performed the analysis for Figure 1, and edited and revised the manuscript. P.D. revised and adapted the script for the deployment of the algorithm in its different forms, set up the release for the web-based application and the python pip package, and edited and revised the manuscript. G.C. acquired funding and edited and revised the manuscript.

## Data Availability

All scripts, tutorials and example images are available at https://github.com/raqueladaia/AxoDen. AxoDen can be used online, as a web application and without the need for installation in https://axoden.streamlit.app/. Additional information can be found in www.corderlab.com and https://sandovalorteqa.com/.

